# ZeptoCTC - Sensitive Protein Analysis of True Single Cell Lysates using Reverse Phase Protein Arrays (RPPA)

**DOI:** 10.1101/2023.09.16.558042

**Authors:** Mahdi Rivandi, André Franken, Liwen Yang, Anna Abramova, Nadia Stamm, Jens Eberhardt, Berthold Gierke, Meike Beer, Tanja Fehm, Dieter Niederacher, Michael Pawlak, Hans Neubauer

**Affiliations:** Department of Obstetrics and Gynecology, Heinrich Heine University of Duesseldorf, Duesseldorf, Germany; Center for Integrated Oncology (CIO Aachen, Bonn, Cologne, Duesseldorf); ALS Automated Lab Solutions GmbH, Jena, Germany; NMI TT GmbH, Protein Profiling, Reutlingen, Germany; NMI Natural and Medical Sciences Institute at the University of Tuebingen, Reutlingen, Germany

## Abstract

Circulating Tumor Cells (CTCs) are commonly analyzed through genomic profiling, which does not capture posttranslational and functional alterations of encoded proteins. To address this limitation, we developed ZeptoCTC, a single-cell protein analysis workflow that combines established technologies for single-cell isolation and sensitive Reverse Phase Protein Array (RPPA) analysis to assess multiple protein expression and activation in individual CTCs. The workflow involves single cell labeling, isolation, lysis, and printing of the true single cell lysates onto a ZeptoChip using a modified micromanipulator CellCelector^TM^. Subsequently, the printed lysates undergo fluorescence immunoassay RPPA protein detection using a ZeptoReader followed by signal quantification with Image J software. ZeptoCTC was successfully optimized, beginning with the measurement of EpCAM protein expression—a standard marker for CTC detection. As expected, mean fluorescence signals for EpCAM levels were significantly higher in single MCF-7 cells compared to MDA-MB-231 cells. Next, Capivasertib-treated MCF-7 cells exhibited an approximately 2-fold increase in the pAkt/Akt ratio compared to non-treated control cells. This finding was consistent with a co-performed western blot analysis of pooled MCF-7 cells. Application of ZeptoCTC to the analysis of single CTCs derived from a metastasized breast cancer (MBC) patient indicated a significantly higher level of pAkt, accompanied by a corresponding increase in pErk level when compared to patient-matched WBC. Finally, the current workflow successfully indicated the detectable pAkt and Akt signal difference in CTCs from two MBC patients: one with an Akt1 wild-type genotype, and the other harboring approximately 80% Akt1(E17K) mutated CTCs. The mutated CTCs revealed clearly elevated pAkt levels (1.8-fold), along with an even more strongly elevated total Akt (3.4-fold) when compared to the respective signals measured in wild-type CTCs. In conclusion, ZeptoCTC is a highly sensitive method for measuring the expression and phosphorylation of treatment-relevant proteins in key cancer-driving signaling pathways from true single cell samples.

## Introduction

Circulating tumor cells (CTCs) are cells released from primary tumors or metastases into the bloodstream, where they are supposed to contribute to further tumor cell dissemination and metastasis. Different methods and technologies have been developed to identify and isolate CTCs (1) from a patient’s blood in order to predict clinical outcomes and indeed the presence of five or more CTCs in 7.5 mL of blood, has been associated with worse overall survival and prognosis in MBC (2).

Apart from their enumeration, uncovering the CTCs’ genotypes and phenotypes has become a major focus in cancer research to understand their biology (3). To this aim, CTCs are isolated and further characterized – due to their stability – mainly at the level of their genomic DNA to uncover DNA mutations and aberrations informing us about potential targeted treatment options (4). Some frequently detected - hence called - hotspot mutations are known for e.g., genes such as *AKT1*, *PIK3CA*, and *ESR1*, reaching frequencies between 2% to 40% in breast cancer (5–7). Such hotspot mutations, which mostly result in over-activated protein products drive the progression of tumor disease and the expansion of metastasizing tumor subclones. Since Alpelisib or Capivasertib drugs to specifically target over-activated *PIK3CA* or *Akt1*, respectively, are available, BC patients might benefit from CTC-based targeted mutation analysis (8–10). However, hotspot mutations (established long-term) may be associated with different protein and pathway activities (short-, mid-and long-term, with high dynamics) – depending on the context of e.g., other mutations in *cis*, therapeutic impetus, or other cross-compensatory posttranslational modifications, or, will not always directly translate into altered protein network perturbations. Therefore, determining the activation of a driver protein or a downstream target protein in single CTCs will provide highly valuable information on the functional range of disease-inducing mutations and will open further – also combined – treatment options (11–15).

The investigation of proteins and – even more important – their activation levels at often low-abundant absolute levels and in single cells is hampered by limited detection sensitivities of currently available protein technologies or their need for high starting cell numbers (16). Such protein analysis approaches encompass highly multiplexed methods including flow cytometry (17), mass spectrometry (18), and mass cytometry–CyTOF (19), or include reports about single-cell absolute protein quantification methods using adapted immunoassay-based Western Blot (20) and ELISA (21). They mostly represent sophisticated, miniaturized versions of mature technologies that reach high sensitivities e.g. through the detection of very dilute (fg/ml) analyte concentrations (22) or added high analyte specificity through the use of NA-based recognition elements (23). However, they are hardly appropriate to analyze single CTCs as starting material since most procedures are not optimized for using single-cell input and suffer from loss of analyte during sample processing (24). Therefore, there is a need for an extremely sensitive protein analysis method, that is integrated into an established and robust single CTC or other single rare cell detection, isolation, and preparation workflow.

Reverse Phase Protein Arrays (RPPA) is a miniaturized, sensitive proteomic immunoassay technology established for focused profiling of multiple signaling proteins and markers in small-sized samples, and has proven to provide valuable quantitative information, especially for the analysis of crude and complex matrices such as cancer cells or tissues. First described by Prof. Lance Liottás group in 2001 (25, 26), RPPA was derived from classical western blot and miniaturized into a ‘dot-blot’-like micro-sample-array format for higher sample throughput, with each sample in a printed array consuming only minute amount of crude sample volume per protein test (nL sample consumption including replicate spots per array, one array = one protein test). With the use of validated specific antibodies for protein analyte detection and up to hundreds of printed replica arrays easily producible from the sample volumes, the approach is capable of quantitatively detecting multiple low-abundant proteins and/or specific protein modifications with high sensitivity. Almost in parallel, Pawlak et al. reported in 2002 (27) on Zeptosens RPPA, adapted as an integrated system solution in combination with planar waveguide technology (27) for ultra-sensitive fluorescence detection, reaching high precision of quantitative signal read-out with the integrated workflow and matching tools and reagents. Over the past years, RPPAs have matured with different versions and workflows and evolved as a protein profiling platform proven to provide sound and valuable information in translational research, drug development, and precision medicine, particularly in the oncology area (28). RPPA measurements with the Zeptosens/ZeptoChip method can detect proteins at low zeptomole levels (27, 29) – hence the name. It detects fractions of low abundant target proteins from an estimated equivalent of a single cell total protein, when printed into one sample array spot from a lysate that is typically prepared from a higher bulk number of starting cells. We have combined this RPPA method with a workflow elaborated to detect and isolate single CTCs and demonstrate for the first time the analysis of signaling proteins from true single cell isolation and sample preparations shown here for single CTC. This work-flow employs (a) the FDA-approved CellSearch® system (Menarini Silicon Biosystems, Bologna, Italy) to identify and isolate CTCs based on their expression of the epithelial cell adhesion molecule (EpCAM), followed by (b) the single CTC isolation, single CTC lysis, and subsequent spotting of single CTC lysates onto hydrophobic ZeptoChips using the state-of-the-art automated micromanipulator CellCelector^TM^ (Sartorius, Jena) (30), and as a last step (c) applying adapted Zeptosens RPPA to process the printed single cell lysate arrays, measure and quantify the signals of performed protein analyte fluorescence assays, and to determine and relate the quantitative signal levels of total and phosphorylated forms of the chosen analyte proteins in the printed samples and controls. We further demonstrate the applicability of this highly sensitive and robust workflow for single-cell protein analysis by determining the relative expression of total and phosphorylated Akt and Erk1/2 signaling proteins in individual CTCs of MBC patients.

## Experimental procedures

### Cell lines

Breast cancer cells from cell lines MDA-MB-231 and MCF-7 were used (ATCC, Manassas, United States; catalog numbers: MDA-MB 231: HTB-26, and MCF7: HTB-22) and cultured in RPMI 1640 containing 10% fetal calf serum and 1% Penicillin-Streptomycin (all Thermo Fisher Scientific) and routinely maintained in humidified atmosphere of 5% CO2 and 95% air at 37 °C. Twenty-four hours before treatment with Capivasertib the culture medium of MCF-7 was replaced by phenol red-free RPMI supplemented with 5% charcoal-stripped FCS. Half of the synchronized were then grown in the above-described medium supplemented with 5 µM Capivasertib, the other half was cultured as before. Cells were negatively tested for mycoplasma.

### Clinical samples

DLA was performed on metastasized breast cancer patients at the Department of Obstetrics and Gynecology of University Hospital Duesseldorf, following an approved protocol (31, 32). Written informed consent was obtained from the patients, and approved by the Ethics Committee of the Medical Faculty of Duesseldorf (Ref-No: 3460) and all procedures were conducted in accordance with the principles outlined in the Helsinki Declaration. Subsequently, DLA samples were cryopreserved and thawed using methods described in a prior study (33).

### Reagents

For single cell lysis, Zeptosens Cell lysis buffer CLB1 (50 mL, Zeptosens #9000) and Zeptosens Cell lysis buffer CSBL1 (50 mL, Zeptosens #9020) were utilized. The Micro pick 48® slide was coated with a solution comprising 2% bovine serum albumin (BSA) in Dulbecco’s phosphate-buffered saline (DPBS). This coating solution was applied to the slide to prevent protein loss during the workflow of printing lysate on the ZeptoChip slide.

### Western blot analysis

RIPA buffer (50 mM TRIS; Sigma-Aldrich, T1503) was used to lyse the cells. After measuring protein concentration with the BCA assay (Thermo Fisher Scientific, 23225), 30 μg of total protein lysate was loaded for each sample onto Mini-PROTEAN® Precast Gels (Bio-Rad, 4568123) with 4 × Laemmli buffer (Bio-Rad, Feldkirchen, Germany, 1610747). Separated proteins were transferred to Immun-Blot® PVDF Membranes (Bio-Rad Laboratories, Inc., Hercules, California). After blocking the membrane with 1% BSA in TBS and 0.1% Tween (Sigma-Aldrich, St. Louis, Missouri) (TBS-T) for 60 min at room temperature, the following antibodies were used: Anti-Akt (pan) (Cell Signaling Technology, 4691) and Anti-phospho-Akt (Ser473) (Cell Signaling Technology, 4060). The membranes were incubated overnight at 4°C. After washing the membranes, they were visualized with horseradish peroxidase-conjugated anti-rabbit IgG (Cell Signaling Technology) at a 1:2000 dilution. Data were analyzed using Image Lab 2.0.1 software and normalized to β-actin expression.

### RPPA

ZeptoCTC RPPA protein analysis was based and further developed on RPPA-established standard routines, equipment, consumables, and reagents using Zeptosens technology as described previously (27) and shortly outlined here. All new ZeptoCTC and RPPA adaptation steps are described separately in the Results section. For standard RPPA, lysates are routinely produced by strong denaturing lysis in CLB1 from bulk cell or tissue sample material. For array printing, the lysates are subsequently diluted ten-fold in CSBL1 to reach final print concentrations of typically 0.2 ug/ul total protein, and printed onto hydrophobic ZeptoChips (6 pre-defined array fields per chip) using routinely a piezo-electric non-contact NanoPlotter2 (GeSiM, Grosserkmannsdorf Germany) with the print samples provided in standard 384-well plates, and printing performed by single droplet deposition (0.4 nl per spot) in duplicate spots per lysate sample. For ZeptoCTC, the whole lysis and printing routine has been newly developed and was transferred to a capillary-based, single cell picking device (CellCelector^TM^ micromanipulator, Sartorius, Jena Germany), modified for in situ lysis and printing as described in Results. Standard cell lysates are routinely co-printed into standard RPPA (MCF7 treated (activated) and control cell lysates, at 0.2 µg/µl protein print concentration in CLB1/CSBL), and were also applied and co-printed into the new ZeptoCTC arrays, as quality control and benchmark. Finally, printed ZeptoChip arrays are blocked with 3% w/v albumin, washed in distilled water, dried, and stored at 4°C in the dark until use.

#### Protein Array Immunoassays

Protein signals are measured in a direct two-step sequential immunoassay using a sensitive and quantitative fluorescence read-out. A single array is probed for each protein marker of interest. The primary antibodies were selected from the current established list of specific RPPA assays (currently 700+). The antibodies are up-front well-characterized for analyte specificity and verified extensively in other studies with various sources of cell and tissue samples. For each protein assay, the primary antibody at respective dilution was incubated in Zeptosens assay buffer overnight (15 h) at room temperature, arrays were washed once in assay buffer and incubated for 45 min with Alexa647-labeled anti-species secondary antibody (Invitrogen, Paisley, UK). Arrays were then washed and imaged in solution using a ZeptoREADER instrument (Zeptosens) using the red laser. Typically, six fluorescence images are recorded for each array at exposure times of between 0.5 and 16 s, including negative control assays in the absence of primary antibody (blank assays) to measure non-specific signal contributions of the secondary antibody reagents. All primary antibodies applied for ZeptoCTC were from Cell Signaling Technology Inc. (Danvers MA, USA): CST 3599 for EpCAM, CST4685 for Akt, CST 4060 for Akt-P-S473, CST 9101 for Erk1/2-P-Thr202/Tyr204.

#### Image Capture and Analysis

For each array (primary antibody and blank assays), the image taken at the longest exposure time without showing any saturation was analyzed using ImageJ software as described in Results in more detail. The mean fluorescence intensity (MFI) of each sample was determined from analyzed mean single spot signals, averaged over the number of blank-corrected, replicate mean single spot signals (typically n=3). Coefficients-of-variation (CVs) were calculated as the ratios of the respective standard deviations (shown as error bars in the graphs) and MFIs.

## Results

### ZeptoCTC workflow

The concept for setting up a true single cell protein analysis workflow was to combine and fully integrate already existing, well-validated sub-workflows of selected mature technology modules (schematically displayed in Figure 1). This includes using the DLA and CellSearch® system to provide well-characterized CTC preparations (A), the CellCelector^TM^ automated micro-manipulation system to isolate single cells by capillary picking from cell culture vessels, to prepare true single-cell lysates and to print *in situ* single cell lysates in array format (B), and the Zeptosens RPPA module for sensitive and robust downstream protein profiling, including the usage of the adapted ZeptoChip platform, matched lysis, and printing buffer reagents, upfront validated specific assay antibodies and customized single cell sample array and data analysis routines (C). The modules of the workflow concept were stepwise realized, adapted, and tested with single cells from tumor cell line cultures and then adjusted afterward to CTC.

**Fig 1.**
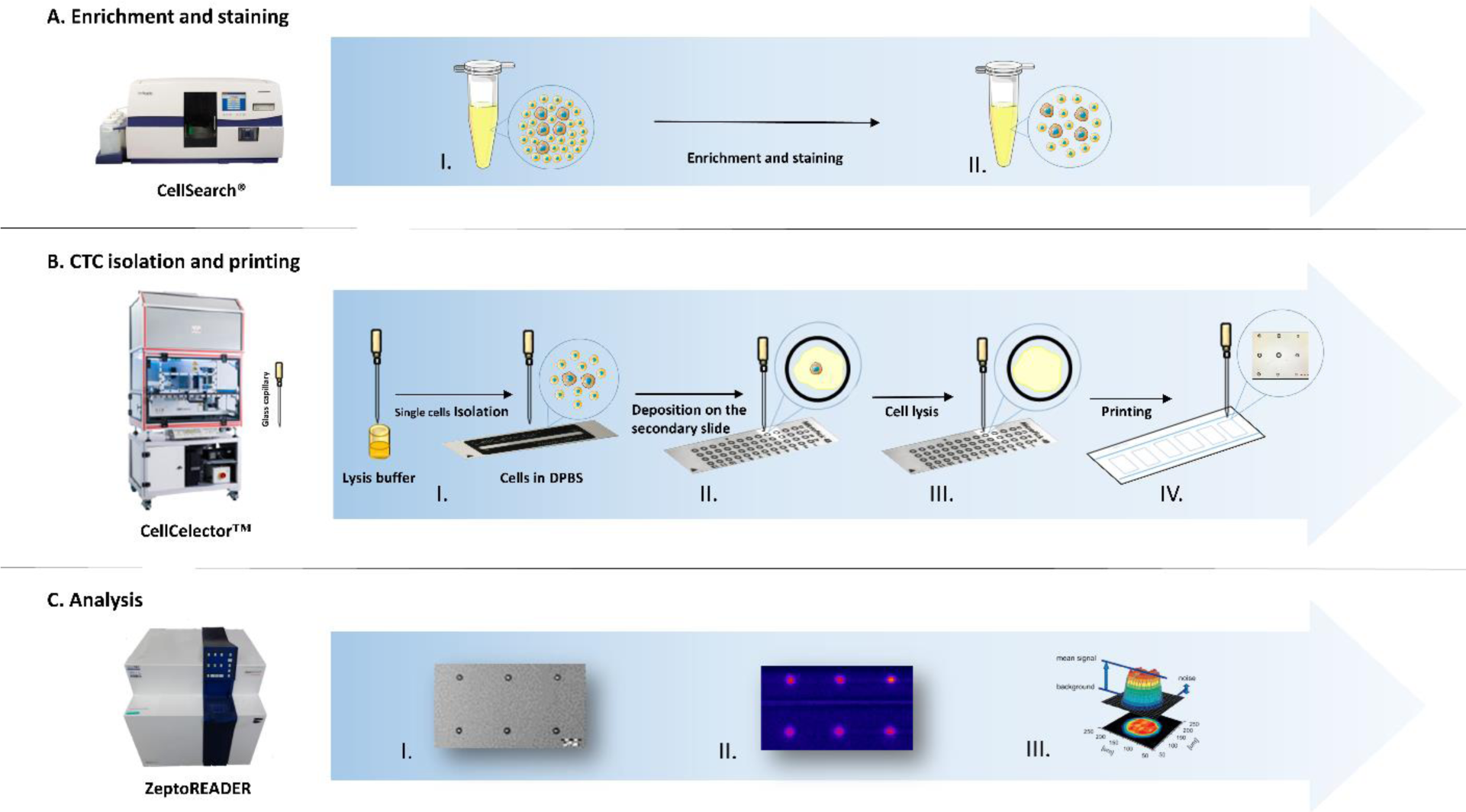
Overview of the established ZeptoCTC workflow. Depicted is the established ZeptoCTC workflow for isolating and analyzing single CTCs with Reverse-Phase Protein Arrays (RPPA). The workflow consists of three primary steps: (A) Capture and verification of CTCs using Diagnostic Leukapheresis (DLA) combined with the CellSearch® system. (B) Single cell/CTC picking with a glass capillary, followed by in situ lysing and mixing on a Micropick 48 slide™ and subsequent printing onto ZeptoChip using a modified CellCelector^TM^ instrument. (C) RPPA analysis of the chip using the ZeptoReader. In panel A, the CTCs are captured using the CellSearch® Profile Kit, and surface markers are stained for imaging. In panel B, a single cell with approximately 20 nL of DPBS is aspirated from a culture dish into the lysis buffer pre-filled glass capillary of the CellCelector^TM^ instrument (total inner volume 40 nL). The CellCelector^TM^ is programmed to deposit 2 nL volume of single cell lysate per sample spot in an array, therefore a total of 40 nL sample lysate volume is sufficient to produce up to 6 sample arrays per chip with 3 technical replicates per sample. Finally, in panel C, downstream RPPA analysis using the ZeptoReader: immunoassays for selected protein markers are performed on the printed and blocked ZeptoChip lysate arrays, fluorescence images of the arrays are recorded and mean spot assay signals quantified with ImageJ software.

It is worth noting that the miniaturization described in this workflow represents a significant improvement over existing standard RPPA techniques. While some RPPA methods, such as Zeptosens RPPA, have demonstrated single-cell sensitivity, they still rely on standard vial 96/384 microtiter plate formats and printing robot routines that require large volumes of buffers with many µL dead volumes, typically necessitating high numbers of starting cancer cells (on the order of 10^4^ to 10^6^). The main challenge of the miniaturized ZeptoCTC approach was to achieve comparably good protein readout performance from true single (or few) cell preparations while ensuring all key specifications for each of the involved technology modules, such as lysis and printing. We performed multiple replicates of lysates obtained from individual cells during the experiments and deposited them onto a chip. This approach enabled us to employ both blank and target assays to measure and quantify the signals, as described in their respective sections or figures. The average fluorescence signals from the printed spots were analyzed using standard Image J software routines (version 1.50f, NIH, USA) and subjected to statistical analysis using a paired t-test. The data analysis was performed using Excel software.

#### Optimizing single cell lysis

A critical step in the workflow was to compose a specific mixture of cell lysis buffer and print buffer to meet the key requirements of this miniaturized one step approach: high efficiency for full cell lysis and good printability from the same composition in parallel; procedures, which for regular RPPA are so far being executed as separate steps (see (29) for more details on buffer requirements). Based on the experience with Zeptosens RPPA lysis buffer CLB1 and print buffer CSBL1, we followed the approach of presenting a concentrated mixture of CLB1:CSBL1 pre-aspirated into the glass capillary and subsequently picking the single tumor cell accompanied with a controlled shot of PBS, which dilutes out the CLB1/CSBL1 mixture. CLB1 has a strong denaturing activity compared to CSBL1 due to its higher urea and thiourea concentrations (7 M urea, 2 M thiourea, and high detergent in CLB1, 3.89 M urea, 1.11 M thiourea in CSBL1, no detergent). On the other hand, CSBL1 containing 11.1% glycerol is beneficial for good printing behavior of the cell lysate and has a low tendency to crystallize. The final buffer composition in a total volume of 40 nL had to guarantee efficient and full lysis at a short duration with high protein yield, prevent crystallization in the glass capillary, and ensure good compatibility with the array printing and final RPPA analysis process. To this aim, both buffers as well as different mixtures of both were selected and tested.

First, we determined completion of cell lysis indicated by changes in the cell’s and nucleus’ morphologies by microscopy in a time-lapse experiment with single cells of cultured MCF-7 breast cancer cell line, which were prelabeled with Celltracker Green CMFDA (Life Technologies, #C2925); final concentration 5 μM (1:2000) and NucBlue™ Live ReadyProbes™ Reagent (Hoechst 33342) (Invitrogen, R37605) in FITC and DAPI and incubated with different CLB1:CSBL1 ratios (1:0, 1:2, 1:5 and 1:10) (Fig 2A). All CLB1:CSBL1 ratios lysed the cells completely in between 1 to 16 min. The fastest cell lysis with approx. 1 min was observed with CLB1-only buffer, followed by the 1:1 mix of CLB1 and CSBL1 with approx. 3 min. Lysis buffer ratios with higher contents of CSBL1 buffer resulted in lysis durations between 7 and 16 min (Fig 2B). After careful consideration, we selected the 1:1 mix of CLB1 and CSBL1, which contained less (50%) detergent than CLB1-only buffer and caused minimal crystallization in the micromanipulator’s capillary. Although CLB1 buffer alone lysed the cells faster than the 1:1 mix of CLB1 and CSBL1, crystallization in the capillary made it unsuitable for our purposes.

**Fig 2.**
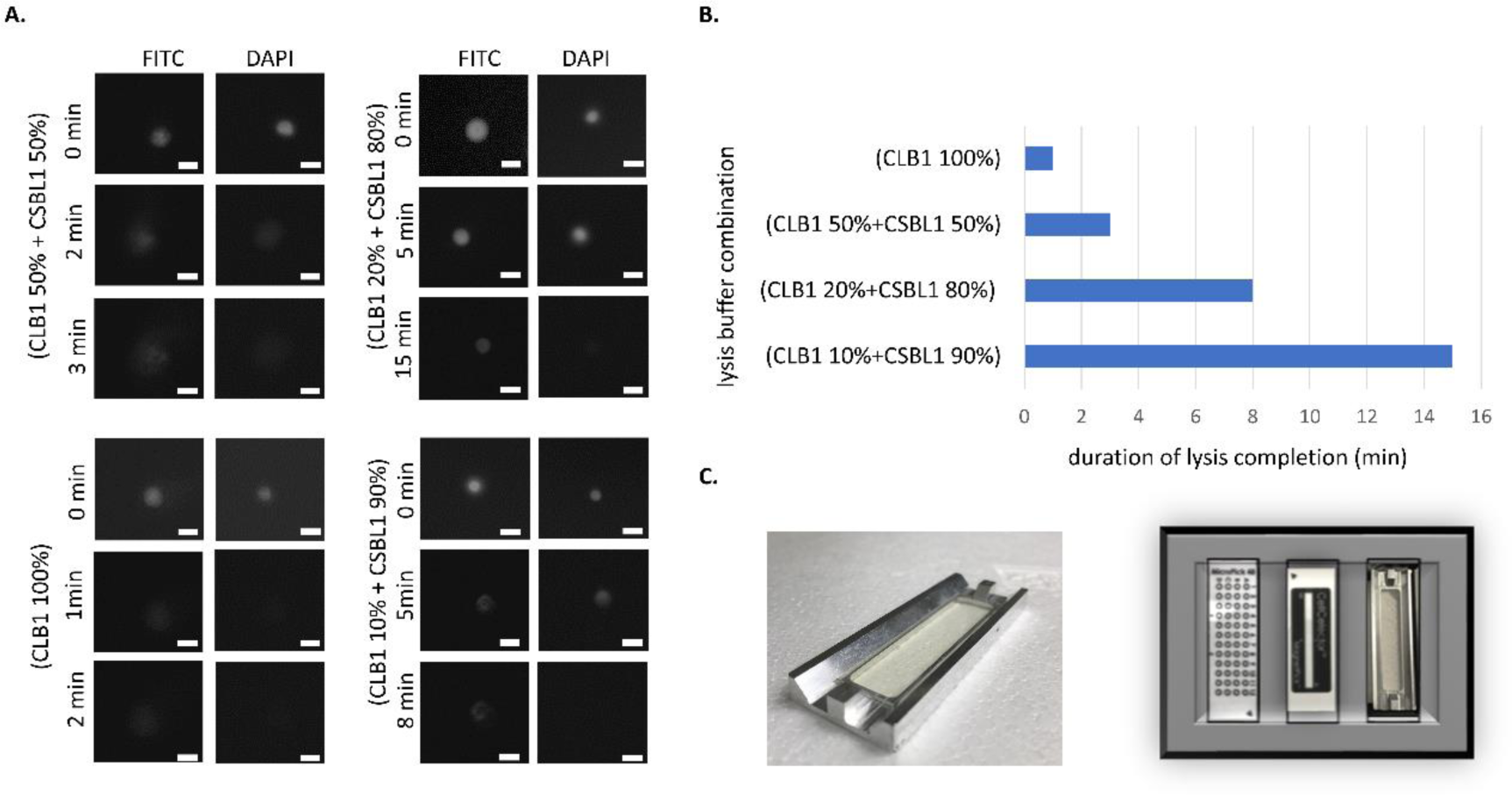
Optimization of conditions and hardware for single cell lysis and manipulation. A) Monitoring of single cell lysis efficiency upon variation of lysis buffer composition. For better visualization, MCF-7 cells were prelabeled with Celltracker Green and NucBlue™ Live ReadyProbes™, and fluorescence images under the microscope were taken at indicated time points during lysis; Scale bar: 25 µm, Magnification:10x; DAPI (nucleus) and FITC (cytoplasm); B) Lysis efficiency: comparison of complete lysis durations (min) for different CLB1:CSBL1 ratios; C) Design of the holder for the ZeptoChip (left) and the adapter for the Micro pick 48® slide, the Magnet pick® slide, and for the ZeptoChip (plus its holder) to position them precisely onto the CellCelector’s microscope stage (right).

In order to optimize single cell lysis and manipulation, additional hardware and software settings were implemented and adjusted. An adaptor was designed and implemented to position the ZeptoChip on the CellCelector^TM^ microscope stage for fast and precise spotting of cell lysates. This setting was highly important to avoid drying artifacts due to the handling of the miniature volumes of single cell lysate (Fig 2C).

Consequently, we finally applied this setting together with CellCelector^TM^ glass capillaries pre-filled with 20 nL of a 1:1 mixture CLB1:CSBL1 and single cells picked with another 20 nL excess volume of DPBS under CellCelector^TM^ control for all subsequent experiments.

#### Single cell lysate printing

To reproducibly print well-mixed single cell lysates with the CellCelector^TM^, a series of steps were taken. Pre-labeled MCF-7 cells were scanned and lysed as described above. Although the lysis procedure immediately started in the micromanipulator’s glass capillary indicating that cell lysis buffer and DPBS were mixed, complete cell lysis and homogenous mixing of all components must be guaranteed. To this aim a total 40 nL of whole single cell lysate was released from the glass capillary into a circled hydrophilic area of a Micropick 48™ slide. The release of the single cell lysate mixture forced instantaneous good mixing of the glass capillary’s content; the completion of cell lysis was video-tracked under the microscope. After a final, efficient mixing by 5 times aspiration-and-release with the glass capillary on the Micropick 48™ slide, the printing procedure of the cell lysate sample started immediately using the same glass capillary and a ZeptoChip adjacently mounted on the CellCelector^TM^ deck (Fig 2C). Technical replicates of single droplet spots (typically n=3) for each sample were printed by aspirating and release of approximately 2 nL volume portions of the complete 40 nL single cell lysate on the ZeptoChip surface, moving the glass capillary to pre-defined spot positions of the 6 pre-defined array fields of the ZeptoChip, one after the other. The lysate volume portions were deposited in close contact with the solution and the ZeptoChip surface. Up to three technical replicate spots per single cell sample were printed as inline series onto each of the 6 array replicate fields per ZeptoChip to examine the reproducibility of printing.

The entire printing process was executed in a controlled atmosphere at 65% humidity. The printed ZeptoChip was kept at RT for 24h followed by a drying incubation for 3.5 hours at 37°C. Finally, ZeptoChip was stored in the dark at 4 °C until further use.

#### Single cell protein analysis

The main difference to conventional RPPA is the way of producing the arrays, i.e., we printed them directly *in situ* after the single cell lysate preparation avoiding dead volumes. The following steps, especially performing the protein assays on the blocked and printed chips were conducted similarly as previously reported (29)

The available array replicates per chip were used to measure in parallel the expression and phosphorylation state of typically 2-4 different protein markers such as the cell surface protein Epithelial Cell Adhesion Molecule (EpCAM) and total or phosphorylated Erk1/2 and Akt proteins –the latter representing the key protein of the PI3K/Akt/mTOR pathway – in a direct fluorescence immunoassay with the proven specific RPPA-tested primary antibodies. The remaining arrays were used as control arrays to address potential auto-fluorescence and blank signal contributions in the absence of specific primary antibodies.

#### Data Analysis of single cell lysate spot signals

The mean signals of the printed spots were analyzed using standard Image J software routines (version 1.50f, NIH, USA) and a flexible array of analysis spot circles. To account for variations in spot size due to capillary-to-capillary or single cell-to-single cell lysate variations, spot signals were analyzed as a mean signal density, with the spot area set constant for all printed spots at a diameter of approximately 150 μm to exclude background pixels. This was well comparable to the 160 μm analysis spot diameter applied in standard RPPA(29) and ensured reproducibility of the quantified assay signals from the technical spot replicates with mean coefficients of variation (CVs) well below 10% with a range between 4% and 7%, see Figure 5A. No significant variation of the mean signals from different arrays was observed. These observations were indicative of the robust printing and complete single-cell lysis processes. Unless otherwise stated, mean spot signals of the primary antibody and the blank assays were averaged to the mean values of the technical replicates (n=2-3). Primary assay means were then corrected for the respective mean blank assay contributions in the absence of primary antibody, but under otherwise comparable conditions and given as Mean Fluorescence Intensity (MFI) values.

### Protein analysis in single cells

#### EpCAM expression

The cell surface protein EpCAM represents the main target to detect and isolate CTCs in MBC. We, therefore, decided to measure EpCAM expression levels first on single BC cell line cells to verify the ZeptoCTC workflow’s performance and to prove the sensitivity of ZeptoChip’s readout. For this purpose, single EpCAM-positive MCF-7 and EpCAM-low/negative MDA-MB-231 cells were isolated and processed as described above.

The lysates of two separately processed pairs of single MCF-7 and MDA-MB 231 cultured cells (cell1, cell2) were printed in technical replicates (2 spots; n=2) onto ZeptoChip replica arrays. On a separate replica array, only a secondary antibody was applied as a blank assay control.

After measuring the signal intensities with the ZeptoReader, the fluorescence images with the longest exposure time (4s) below pixel saturation (16 bit) were analyzed and single cell lysate spot signals were quantified as blank-corrected mean spot signals averaged over the technical replicates. As outlined in Fig. 3, the EpCAM MFI signals for both MCF7 single cell preparations were detected well above blank, and were – as expected – significantly higher than for the EpCAM-low expressing MDA-MB 231 cells. The MDA-MB-231 cell 1 revealed very low, but still detectable residual EpCAM expression levels, for cell 2 no signal was detected. Moreover, these first workflow experiments yielded <10% mean signal CVs ranging from 3% to 8% of the printed technical spot replicates, and hence indicated a good spot-to-spot reproducibility of cellular EpCAM protein detection (see also low error bars in Fig. 3 indicating standard deviations). Additionally, the EpCAM signals of two cells randomly picked from the same cell line (MCF7, MDA-MB-231) were considerably different pointing towards well-known cell-to-cell heterogeneity. Nonetheless, such single cell protein variability and environmental changes seem now addressable with the presented method.

**Fig 3.**
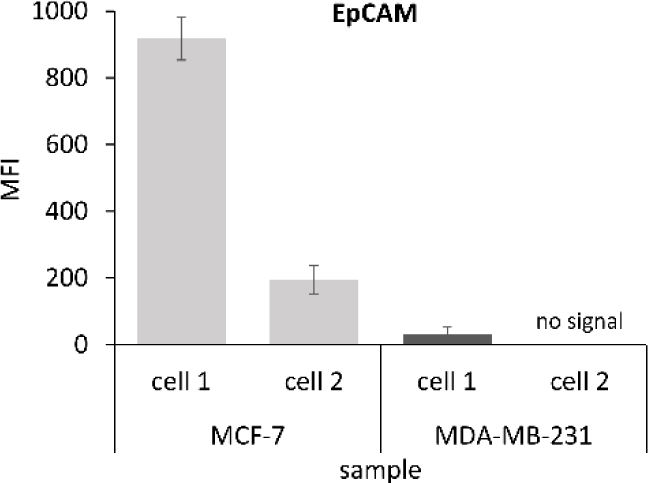
EpCAM protein expression signals measured from single cell preparations using ZeptoChip. Quantification of fluorescence signals depicted from two MCF-7 and two MDA-MB-231 single cell lysate samples, respectively. The mean fluorescence signals (MFI) of both MCF-7 single cells were significantly higher than those of both MDA-MB-231 cells (p-value = 0.0435). Error bars indicate the standard deviations of the mean technical replicate spot signals (n=3).

#### Measuring phosphorylation of Akt in single tumor cell line cells upon Capivasertib treatment

In the next step, we tested if our workflow can be used to measure in parallel expression level and its phosphorylation status of a signal protein of interest. Since the PIK3CA/Akt signaling pathway represents a key pathway not only in BC development but also in its therapy resistance, we selected the Akt protein and its functional form phosphorylated at serine 473 (Akt-P-Ser473), in short pAkt, as key analytes. Phosphorylated Akt was also good to proof for sensitivity, since especially this phosphorylated analyte (and also others) is known for its relatively low abundance and hence a challenge of detection. MCF-7 cells were treated with 5 µM capivasertib for 24 h. Capivasertib is an Akt1-specific inhibitor and is known to lead to elevated pAkt levels, due to its ATP-competitive mechanism of action and accumulation of inactivated hyper-phosphorylated Akt (34). The increased pAkt levels were verified by Western Blot analysis of lysates from bulk MCF-7 cells (Fig. 4B). From the same cell culture used for Western Blot analysis, capivasertib-treated single MCF-7 cells (MCF-7/treated, 3 single cells) and untreated control cells (MCF-7/untreated, 2 single cells) were micromanipulated and processed according to the established ZeptoCTC workflow. Four technical replicates per single cell lysate were printed for each replica array. The arrays were incubated with primary antibodies against Akt and pAkt, followed by the anti-species Alexa Fluor™ 647 conjugated secondary antibody and fluorescence detection with the ZeptoReader (Fig. 4). One array was used as blank control.

**Fig 4.**
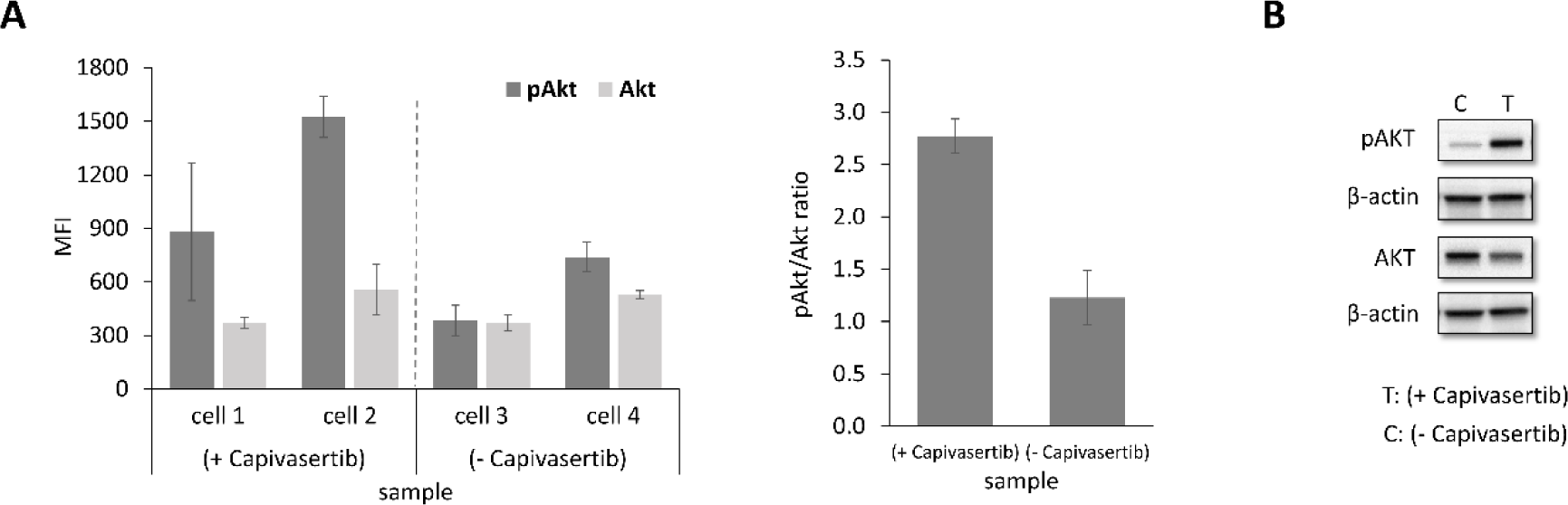
ZeptoCTC expression signals of Akt and pAkt (Ser473) in treated single MCF-7 cells. A) (left) RPPA mean signals (MFI) representing Akt and pAkt levels, and (right) calculated pAkt/Akt mean signal ratios as measured in picked MCF-7 single cells treated with capivasertib (+) and without this treatment (-) as a control using the established ZeptoCTC workflow; result: significantly higher phosphorylation of Akt in treated over non-treated single cells B) Western Blot verification of elevated pAkt from co-performed bulk cell pellet lysate preparations (>2 106 cells) of MCF7 cell line cultures treated with capivasertib and without this treatment as a control.

Blank-corrected MFI fluorescence signals for pAkt in single MCF cells were significantly higher compared to the total Akt protein (see Fig. 4). Treatment-to-Control Ratio (TCR) for pAkt as calculated from the respective single cell MFI mean values showed clear and significant up-regulation (2.4, p = 0.0047) in MCF7-treated over MCF7-non-treated single cells (Fig. 4A left). In contrast, the TCR of total Akt protein remained almost unchanged (1.1), as expected from capivasertib’s exclusive effect only on the functional phosphorylation of Akt (see Fig. 4A left). Consequently, the mean TCR of the Akt-normalized pAkt/Akt ratios was then also more than 2-fold significantly higher (2.8, p = 0.0012) in the MCF7/treated over the MCF7/non-treated cells (see Fig. 4A right). The observed single cell results have been verified by co-performed Western Blot analysis with bulk cell pellet lysate preparations (>2×10^6^ cells starting material) from respective same treated and non-treated MCF7 cell line cultures, (with about 3-fold TCR of pAkt/Akt (see Fig. 4B).

In summary, the data obtained with ZeptoChip detection and true single cell lysates were well in accordance with western blot analysis measured from more than 2×10^6^-fold starting cell material and support that the established workflow is highly sensitive and able to measure both the expression and functional (phosphorylation) status of signaling protein markers in parallel from single cells and in a multiplex format.

### Application of the single cell protein analysis workflow to CTCs

In order to adapt the workflow to CTCs, we decided to use fresh frozen aliquots of DLA-products for mainly two reasons (32). First, the cells are not treated with a fixative such as paraformaldehyde, keeping the CTCs viable and avoiding negative effects hampering cell processing and the measurement of proteins and their phosphorylation levels. Second, since one aliquot is always inspected with the CellSearch® System, both CTC numbers and their expected quality were known. It was also essential to prevent the permeabilization of the cell membrane to avoid the loss of intracellular proteins.

#### Detection and processing of single CTCs

DLA aliquots were processed with the CellSearch® profile kit and the CellSearch® system to capture CTCs, which were identified with a mixture of antibodies targeting only surface proteins. Selection of the used antibodies, their type as well as applied detection labels were carefully coordinated with the RPPA detection routines. The labeled cells were scanned with the CellCelector^TM^ automated micromanipulator (Sartorius, Jena), and single CTCs were isolated and lysed, as described before for the single cancer cell preparations.

The isolated single CTC lysates were printed onto the ZeptoChip slide in the array formats as described before.

#### Measuring phosphorylation of Akt and Erk1/2 in CTCs derived from a MBC patient

Since we aimed to measure protein expression level and in parallel its functional phosphorylation status in single CTCs, we selected a cryopreserved and CTC-positive DLA product obtained from an MBC patient (patient 1). Prior DNA sequencing analysis had confirmed that approx. 80% of her CTCs harbored the activating Akt1(E17K) mutation (35).

Although this mutation should result in the phosphorylation of the Akt protein at serine 473(36), we did not know at which level and frequency. This sparked us to also measure the phosphorylation level of Erk1/2 protein. This protein is a key regulator of the MAPKinase signaling pathway for cellular growth, often interacting directly with the PI3K/Akt pathway in a cross-compensatory manner. For total Erk1/2 and phosphorylated Erk1/2 (Erk1/2-P-Thr202/Tyr204) proteins, well validated and measurable assays were available (since these protein forms are often presented at higher abundance/active state than e.g., pAkt), as part of our established list of RPPA assays (700+ antibodies). For this experiment, we also included – apart from the single CTC samples - lysates of single patient-matched WBCs, since in contrast to treated or untreated cell lines we do not dispose of ‘unstimulated’ CTCs from the same patient. Furthermore, we included validated RPPA standard lysates, which we regularly apply in routine RPPA protein profiling studies, as quality control (QC) for array printing and assay performance. These standard QC lysates had been prepared upfront and characterized from large bulk amounts of MCF7 tumor cell line cultures (>10×10^6^ cells). These standard samples came as aliquoted pairs of treated and control (non-treated) MCF7 lysates (kindly provided by NMI TT Pharma services, Reutlingen, Germany) with pre-confirmed Akt and pAkt (and other protein) levels, and were co-printed onto the ZeptoCTC replica arrays of this experiment, at a concentration representing also single cell equivalent total protein material per spot, together with the true single CTC and WBC sample lysates.

Two different CTCs were isolated from a thawed DLA product and processed as described above (Fig. 5A). All CTCs, WBCs, and standard cell lysates were printed in technical replicates into ZeptoChip replica arrays, with single cell lysate volumes sufficient to determine the multiple Akt-P-Ser473 (pAkt), Erk1/2-P-Thr202/Tyr204 (pErk) protein and blank assay signals, respectively. The quality of the printed arrays is illustrated in Figure 5 (top) with clippings of the fluorescence image recordings. The lysate print achieved homogeneous spot morphology, uniform spot diameters and a good signal reproducibility of the printed spots, evident from a low mean coefficient of variation (CV = 6%) of the sample MFIs quantified by the ImageJ software and averaged overall the different samples printed on the chip.

**Fig 5.**
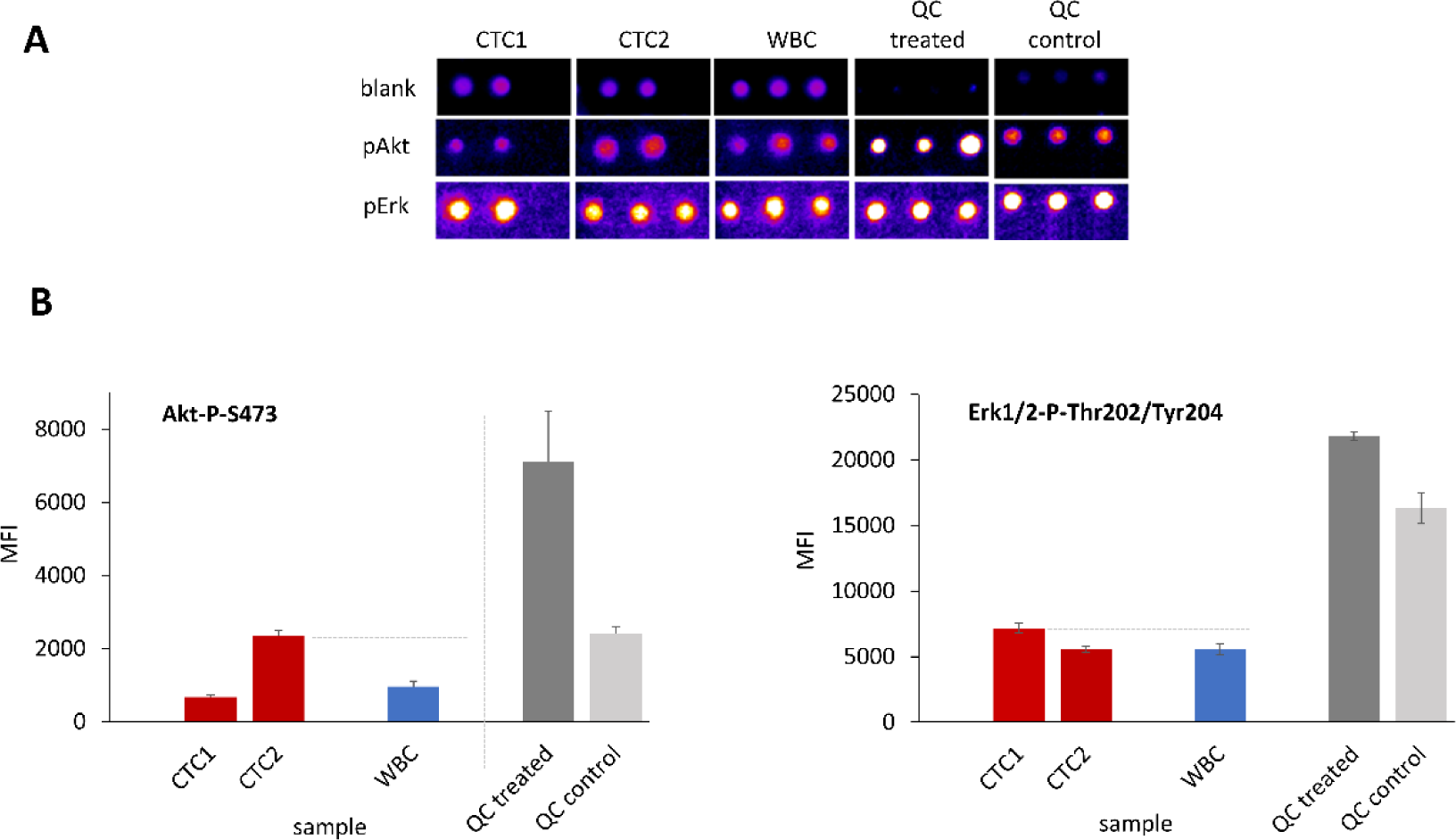
Evaluation of phosphorylated Akt and Erk1/2 proteins in single CTCs derived from a MBC patient. A) False color clippings of RPPA Zeptosens fluorescence array images (exposure 4s) for analysis of pAkt and pErk1/2 protein levels in CTC and WBC single cells from a MBC patient, and comparison to bulk cell QC (MCF) controls (3 replicate spots printed per sample). B) Mean fluorescence signals (MFI) as quantified by ImageJ: elevated pAkt and pErk1/2 levels in CTCs compared to WBCs in line with QC lysates; pErk1/2 abundancy signal 3-11 times higher than pAkt as evident from different MFI scales.

The blank-corrected mean fluorescence intensities (MFIs) showed more than two-fold difference among the different CTCs and were clearly more pronounced for pAkt than for pErk1/2 levels (see Fig. 5B). One of the two printed single CTC lysates (CTC2) showed a clear and significantly higher pAkt level compared to the patient-matched WBC (2.4-fold, p=0.02), whereas the pErk1/2 levels (CTC1 and CTC2) were almost comparable (1.2-fold, p=0.02). Notably, the pErk1/2 abundancies were 3-11 times higher than for pAkt as evident from the different absolute MFI scales (Fig. 5B). Significantly elevated pAkt and pErk1/2 levels observed in the treated standard QC lysates were well in agreement with the measured enhanced signals in the CTCs. Also, the absolute signal levels of the co-printed standard QC controls and the single CTC lysate MFIs were well in a comparable order range, even though they were prepared according to completely different protocols (one prepared from bulk cells and the other from only single cell amounts). Besides, the treatment-to-controls (TCR) ratios of the standard QC lysates were at well comparable values and quality when printed and measured (i) in a routine RPPA setting (> 10E6 cells lysed at ml volumes) and (ii) in the new, true single cell lysate workflow (single cells lysed at few nL volumes).

These data demonstrate that the new ZeptoCTC workflow can measure multiple proteins expressed with low and high abundancies from true single cell sample preparations.

#### Measuring phosphorylation of Akt protein in single CTCs derived from two MBC patients

As a next step, we aimed to investigate the potential of the ZeptoCTC workflow in measuring expression and activation signals in CTCs derived from patient samples, both with and without the Akt1(E17K) mutation. To achieve this purpose, cryopreserved and CTC-positive DLA products were obtained from patient 1 [Akt1(E17K)] and another MBC patient (patient 2) with CTCs of wild-type *Akt1* genotype (Supplement Seq Data). Single CTCs and WBCs were picked with the CellCelector^TM^ and processed as described above (see Fig. 6A and 6B for results). The printed single cell CTC lysates of the two patients, co-printed patient-matched WBC lysates, and standard QC lysates were investigated, this time with a focus on pAkt and total Akt protein levels. As indicated by the low standard deviations and the QC standard lysate signals (Fig. 6), the quality of the printed arrays and the reproducibility of the replica spot MFI assay signals were again good and successfully attained. The CVs of mean replica spot MFI assay signals averaged over all printed CTC and WBC single cell lysates of patient 1, were 3% for the pAkt and for the total Akt. For the CTC and WBC single cell lysates of patient 2, similar low mean CVs were achieved (5% for pAkt and 3% for total Akt, respectively). With these numbers, print and assay signal CVs of the ZeptoCTC application were well comparable with those reported for standard RPPA applications (see (29) for numbers). The MFI signals and TCR of the co-printed standard QC lysates for both pAkt and Akt confirmed the correct readout process for both pAkt and total Akt. In the CTC single cells from patient 2 with AKT wildtype CTCs, both the Akt and pAkt average absolute signals were slightly below than in the patient-matched WBC (0.9-fold for pAkt, 0.8-fold for Akt), see Fig. 6B left. This resulted in about comparable pAkt/Akt mean signal ratios (5.4 for CTC, 4.1 for WBC), see Fig. 5B right. The pAkt/Akt signal ratios indicate that also pAkt/Akt levels in these two cell types were almost comparable. In CTCs isolated from patient 1 with AKT1 mutated CTCs, slightly higher absolute pAkt (1.1-fold) and almost double-fold Akt signals (1.9-fold) were observed compared to the respective WBCs (Fig 6A left). This resulted in reduced pAkt/Akt signal ratios in the CTCs (mean 2.4-fold) compared to the WBCs (mean 4.5-fold, comparable to the respective wild-type ratio), see Fig. 6A right, mainly due to increased relative total Akt in the mutated compared to wild-type CTCs.

**Fig 6.**
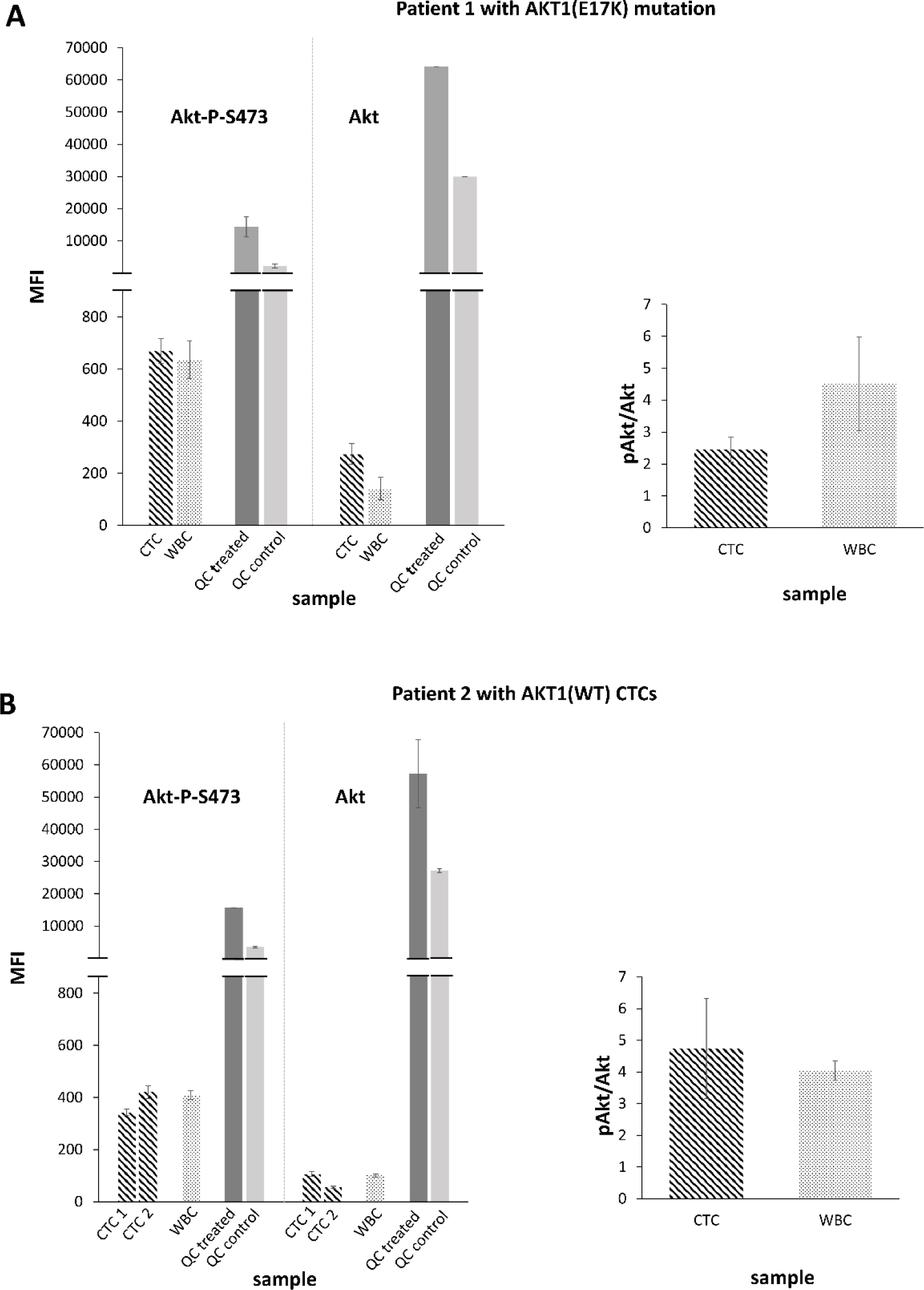
Measuring phosphorylation of Akt to Akt protein in single CTCs derived from two index metastasized breast cancer patients. A) (left) RPPA Zeptosens analysis of pAkt and Akt protein signals (MFI) in single CTC and WBC samples derived from index patient 1 harboring an Akt1(E17K) mutation, in addition to co-printed quality control samples (QC treated, QC control); (right) pAkt/Akt mean signal ratios of respective patient 1 single CTC and WBC samples. B) (left) RPPA Zeptosens analysis of pAkt and Akt protein signals (MFI) in single CTCs and WBCs from index patient 2 (wild type); (right) pAkt/Akt mean signal ratios of respective patient 2 single CTC and WBC samples.

Assessing also the good reproducibility of the quality control signal intensities in both chips (mean CV < 10%), in the patient with harboring CTCs with Akt1(E17K) mutation, the absolute MFIs of the CTCs lysates in the pAkt assay were clearly higher (1.8-fold) than those of the CTC lysates from the patient with Akt1(WT) CTCs (670 ± 47 for mutated versus 382 ± 39 for wildtype), and were even higher (3.4 fold) in the total Akt assay (274 ± 39 for mutated versus 81 ± 26 for wild-type). Although we do not know the Akt genotype of each CTC investigated, this observation suggests that the ZeptoCTC workflow has the capacity to measure a higher signal intensity for pAkt in CTCs obtained from the MBC patient with Akt1(E17K) mutated CTCs compared CTCs from the MBC with Akt1(WT) CTCs.

In summary and in line with our quality control measures, these data demonstrate that ZeptoCTC is a robust workflow and able to measure target proteins in single CTCs. Furthermore, the possibility to process up to 6 replica arrays/assays in a single chip run with a number of true single cell lysate preparations, opens the window to analyse several key pathway protein markers in parallel in a flexible (RPPA like) multiple assay and sample setting.

## Discussion

This proof-of-concept study has successfully developed ZeptoCTC, an ultra-sensitive protein analysis tool for single cells, including CTCs. ZeptoCTC integrates standardized and highly sensitive systems to enrich, detect, and isolate single CTCs and to measure low levels of on and off-target proteins and their phosphorylation status. With this breakthrough, we anticipate that ZeptoCTC will effectively link single-cell DNA genotype information to its functional outcomes, such as tumor growth and resistance to therapy. This integration will greatly enhance precision medicine, bringing us one step closer to improved treatment outcomes.

In the current landscape, molecular analysis of CTCs is a useful tool to identify and monitor targeted treatment options, especially when tissue biopsies are unfeasible. However, genetic changes in nucleic acid sequences may not always provide a complete picture of protein products and especially no information about their activation status, such as phosphorylation levels or subsequent signaling pathway activation. Therefore, understanding the gene-phenotype association in single tumor cells is crucial. Protein activity is regulated by various cellular contexts, including other genetic mutations and post-translational modifications. Empirical evidence from BC cell lines has shown that the response to PI3K inhibitor Alpelisib, despite harboring a hotspot PIK3CA mutation, overlaps with PIK3CA wild-type genotype counterparts (11). Moreover, sequencing methods have revealed multiple tail mutations located outside of hotspot regions (37). For instance, certain double mutations in cis to hotspot mutations have been documented to enhance PI3K activities, downstream signaling, cell proliferation, and tumor growth compared to ‘only-single-hotspot-mutated’ PIK3CA BC, whereas others exerted so far unknown effects on protein activities (14, 38–41). Similarly, the Akt gene showcases similar dynamics, where three out of six novel pleckstrin homology domain mutants (L52R, C77F, and Q79K) predominantly found in human BCs are activating oncogenic mutations (42). These examples underscore the need for comprehensive functional testing of cancer gene variants, ideally conducted directly in isolated tumor cells.

Understanding single-cell protein analysis is not only fundamental for unraveling cell biology but also for identifying patients who can benefit from targeted therapies. When combined with DNA mutation data, a single CTC protein analysis can offer vital additional insights for therapy selection and accurate disease state determination.

### Key results

The study’s key results emphasize ZeptoCTC’s capability to consistently generate well measurable and valuable multi-protein expression data at the true single-cell level by applying a more miniaturized, sensitive RPPA read-out. This includes good signal-to-noise ratios and low signal variations with CVs < 10% among the technical replicate spots printed from the prepared true single cell sample lysates. Furthermore, a low variation was observed between processed chips, as evident from reproducibly measured pAkt and Akt signals obtained from co-printed, up-front validated RPPA control lysates.

Due to this precise performance of ZeptoCTC, we argue that the observed signal variabilities observed between single cells – also of the same cell line – are pointing toward well-known cell-to-cell heterogeneity, a phenomenon that has been previously noted in other single cell experiments (43–45). For instance, EpCAM expression levels in two MCF7 cells were highly different, matching observations we have made previously suggesting a continuum of EpCAM-positivity in MCF-7 cells (46). Similarly, varying levels of Akt and pAkt in single MCF7 cells treated with Capivasertib suggest diverse drug responses within individual cells. In addition, ZeptoCTC demonstrated its high sensitivity and reproducibility by enabling the detection of signals for the putatively low-expressed Akt protein in its total and phosphorylated form (Akt-P-S473), even when (currently) printed from only a 1/18 volume of the complete prepared single cell lysate per spot. This opens up the possibility of designing ZeptoCTC assays that cover more proteins of other key signaling pathways or multiple (like a handful) number of other key protein markers for more comprehensive investigations. Specifically, we envision an assay that scrutinizes all proteins currently targeted in BC involved in the Her2-PIK3CA-Akt1-mTOR signaling pathway. To achieve this goal, further titration experiments will be conducted.

### ZeptoCTC’s advantages and success in detecting low abundance proteins

ZeptoCTC holds a unique advantage over other published single cell protein analysis tools as it combines various methods that have individually demonstrated suitability for single cell identification, processing, and protein analysis with high sensitivity. This on-purpose-built application begins with the unequivocal recognition and separation of a pure tumor cell, which minimizes confounding background and results in minimal loss of cells and cell lysate. Theoretically, ZeptoCTC enables to analyze of a blood sample containing only one CTC in a highly specific and sensitive manner.

The success of ZeptoCTC in detecting low-abundance proteins in single cells is attributed to its use of highly specific antibodies for both CTC detection and protein analysis and its high physical readout sensitivity based on planar waveguide-based fluorescence detection. This allows the assay to achieve a demonstrated limit of analyte protein detection in the zeptomole range, i.e., the detection of a signal from roughly 1000 protein molecules (∼0.1 fg) in one spot of total of 10^9^ protein molecules (∼100 pg) through the bench-top ZeptoREADER. The study’s high assay sensitivity was evident in the present study, where the mean fluorescence intensity of MCF7 single cell triplicate lysate droplets was higher than MDA-MB 231 cell lysate regarding EpCAM expression. Further enhancing assay sensitivity, the sample lysis volume was adjusted to approximately 40 nL for every single cell, with only 2 nL print volume applied per replicate spots on the ZeptoChip surface, leading to less single cell sample lysate consumed per array and demonstrating higher sensitivity compared to other methods such as SCoPE-MS(47) and OAD and nanoPOTs(48, 49), which have larger sample processing volumes.

### Advantages of the approach and enrichment of patient CTCs

Our approach, which involves lysing and printing a targeted single cell, offers a crucial advantage over other methods that utilize bulk proteome digestion to estimate the theoretical protein amount of a single cell. This advantage becomes particularly relevant in translating ZeptoCTC to the analysis of CTCs in patient blood, where tumor cells are present in limited numbers. The use of bulk lysate approaches in such cases can introduce technical and biological confounding factors since analytical frameworks developed for bulk methods are not necessarily suitable for single-cell analyses.

In our assay, the reference quality control lysate, prepared from a bulk of treated and control cells, was scaled down to a single cell, and despite showing a significant difference between the treated and control groups, revealed much higher mean fluorescence signals compared to single CTC and WBC.

In the realm of single-cell proteomics analysis of CTCs, only a handful of studies have utilized real single CTCs from patient blood to detect their low abundance of proteomic content (50, 51).

To address this challenge, we developed a comprehensive workflow that enables the enrichment, isolation, and printing of lysates from patient CTCs.

Our optimized antibody cocktail, when used with the CellSearch® platform, enabled us to enrich and isolate rare CTCs with high precision using the CellCelector^TM^. Importantly, our workflow is not limited to specific enrichment methods and can potentially be adapted for use with other approaches.

In another avenue of research, a distinct study employed a combination of FACS and the nanoPOTS platform, bolstered by automated LC-MS systems, to delve into single-cell proteomics. This effort led to the identification of 252 proteins (49). However, it’s essential to underscore that this particular study did not extend its analysis to encompass real single CTCs. In contrast, our innovative approach harnesses the capabilities of an embedded live camera within the CellCelector^TM^ device. This technological innovation introduces real-time cell imaging, revolutionizing the entire process by substantially enhancing the scrutiny of each step’s efficacy and spanning the isolation, lysing, and printing of cells.

### Limitations and Outlook

Despite the well reproducible results obtained from every triplicate of single cell lysate and the high sensitivity in detecting targeted proteins in real CTCs, several limitations of the current ZeptoCTC should be acknowledged. These include mainly technical limitations, like the time-consuming nature of the process with the current instrumentation which is not yet fully automated and integrated to facilitate an easy, flexible, and faster printing of higher sample numbers, or, challenges in software analyzing irregular spot morphologies, and handling variations in spot size and blank corrections. To address these limitations, the current state developed workflow is flexible and allows for the integration of alternative printing techniques and software solutions that can enhance both printing and assay sensitivity. While the current workflow has been successful in detecting low abundance protein modifications at the real CTC level, it is still limited in terms of multiplicity, both in terms of the number of CTCs and the targeted proteins. Looking ahead, modified chips with a greater number of arrays can be developed to enable the analysis of important molecules in targeted pathways. This will be essential for achieving more precise personalized therapy. And, our approach is adaptable to other rare single cell applications.

Another limitation we encounter in our study is the absence of simultaneous analysis of both DNA and protein within the same circulating tumor cells (CTCs). This has hindered our ability to definitively ascertain the presence of specific genetic mutations in the CTCs that underwent protein analysis. Upon genetic evaluation of a subset of CTCs, we discovered that roughly 20% of these cells did not exhibit the anticipated mutation. Consequently, the potential exists for inadvertent selection of CTCs with normal genetic profiles, rather than those with the intended mutation. To address this, a potential avenue could involve developing a technique for isolating the cell nucleus and segregating the cytoplasm, thus enabling more accurate protein analysis. Additionally, leveraging specific antibodies tailored to detect particular mutations, such as those affecting the AKT gene, could aid in identifying CTCs harboring the targeted mutations. Implementing these enhancements could substantially enhance the accuracy and reliability of our exploration into the intricate interplay between genetic mutations and protein expression within the CTC context.

## Supplemental data

This article contains supplemental data.

## Conflicts of interest

The authors declare that they have no known competing financial interests or personal relationships that could have appeared to influence the work reported in this paper.

## Acknowledgments

This project was funded by the Deutsche Krebshilfe (German Cancer Aid; #70112504). Figure 1 was partly generated using Servier Medical Art, provided by Servier, licensed under a Creative Commons Attribution 3.0 unported license.

## Author contributions

Hans Neubauer, Michael Pawlak: Supervision, Conceptualization and Methodology; Mahdi Rivandi, Anna Abramova, Nadia Stamm: Data curation, conducted experiments; Mahdi Rivandi, Liwen Yang: Collated clinical tissues; Berthold Gierke, Meike Beer, Mahdi Rivandi: Visualization, Software; Mahdi Rivandi, Hans Neubauer, Michael Pawlak, André Franken: wrote the manuscript; Tanja Fehm, Jens Eberhardt, Dieter Niederacher: Resources, Reviewing and Editing.

## Abbreviations

Bovine serum albumin: BSA
Breast Cancer: BC
Cell Lysis Buffer: CLB1
Cell Spotting Buffer Lysate: CSBL1
Circulating tumor cell: CTC
CyTOF: Cytometry by the time of flight
Diagnostic leukapheresis: DLA
Dulbecco’s Phosphate Buffered Saline: DPBS
Epithelial Cell Adhesion Molecule: EpCAM
Mitogen-activated protein kinase: Erk1/2
Estrogen Receptor 1: ESR1
Half-maximal inhibitory concentration: IC50
Metastatic breast cancer: MBC
Mean Fluorescence Intensity: MFI
Nucleic acids: NAs
phosphorylated Akt: Akt-P-Ser473 or short pAkt
phosphorylated Erk1/2: Erk1/2-P-Thr202/Tyr204 or short pErk
Phosphatidylinositol-4,5-Bisphosphate 3-Kinase Catalytic Subunit Alpha: PIK3CA
Phosphoinositide-3-kinase: PI3K
Primary tumor: PT
Protein kinase B: Akt
Reverse Phase Protein Arrays: RPPA
White Blood Cell: WBC

## Notes

### Competing Interest Statement

The authors have declared no competing interest.

