## Supplementary Materials for "ZeptoCTC - Sensitive Protein Analysis of True Single Cell Lysates using Reverse Phase Protein Arrays (RPPA)"

Table S1. Clinicopathological characteristics of the two metastatic breast cancer patients

Video S1: On-stage demonstration of the established workflow, featuring the following steps: 1. Deposition of isolated single Circulating Tumor Cells (CTCs) on a Micropick 48 slide™, 2. Lysing the isolated cell on the slide, and 3. Printing the lysate onto ZeptoChip slides

Figure S1. Mutation analysis of the AKT1 E17K hotspot in two index breast cancer patients.

Figure S2. Fluorescence microscopy of two CTC and WBC from a breast cancer patient using CellCelector^TM^ stage microscopy.

**Table S1. Clinicopathological characteristics of the index metastatic breast cancer patients.**

| Patient ID | Age | Tumor size* | Nodal status* | Metastasis status* | Histology | Grading | Molecular Subtype | Lines of therapy in metastatic situation |
| --- | --- | --- | --- | --- | --- | --- | --- | --- |
| AKT1 mutated | 74 | 2 | 0 | 0 | Invasive lobular | 2 | Luminal | 2 |
| Not AKT1 mutated | 46 | 1 | 1 | 0 | NST | na | Luminal | 1 |

*At the time of diagnosis; na: Not analyzed

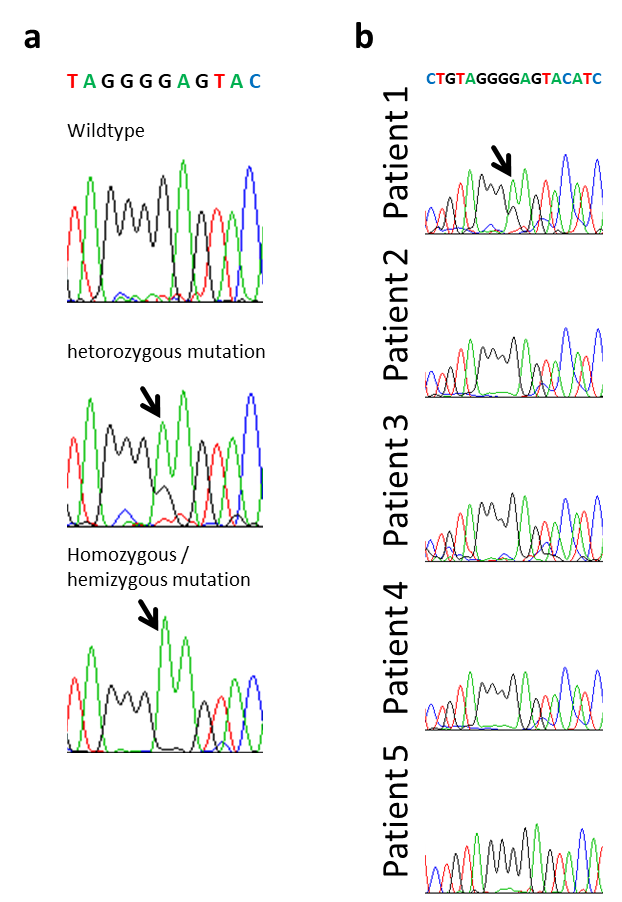

A

B

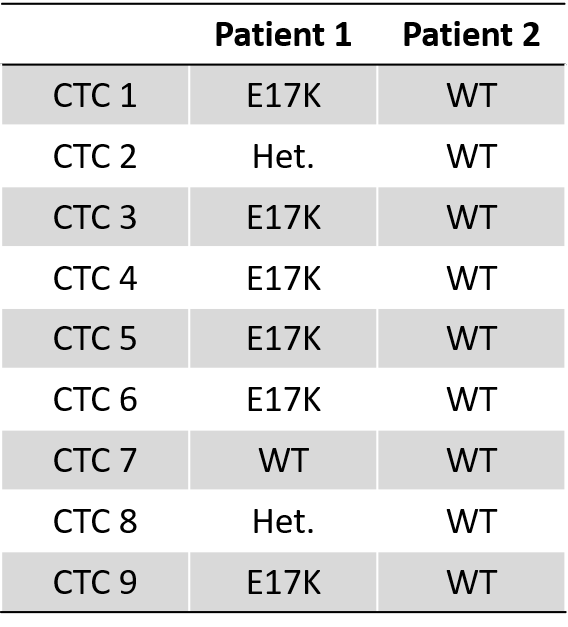

**Figure S1. Mutation analysis of the AKT1 E17K hotspot in two index breast cancer patients.**

(A) References for E17 hotspot mutation in AKT1. The Analysis was performed by Sanger Sequencing. (B) The E17 hotspot region of AKT1 was analyzed on WGA products from 9 CTCs of each patient. WT, wild-type; Het., heterozygous

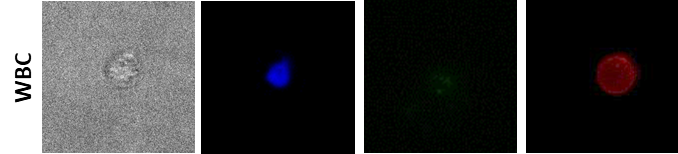

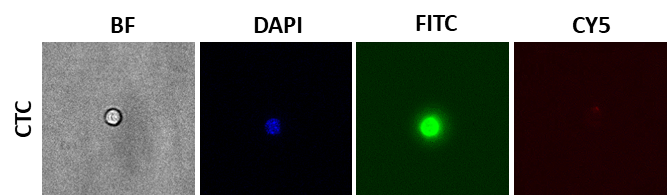

50 µM

25 µM

**Figure S2. Fluorescence microscopy of two CTC and WBC from a breast cancer patient using CellCelector^TM^ stage microscopy.**

In Panel A (CTC), CTC exhibits surface markers' signals for cytokeratin (FITC) and nuclei (DAPI) while demonstrating a negative signal for CD45 (CY5). The scale bar in Panel A measures 50 µM. In Panel B (WBC), WBC presents nuclei (DAPI) and CD45 (CY5) signals, with a negative signal for cytokeratin (FITC). The scale bar in Panel B measures 25 µM.
